# The N-terminus of Sec61p plays key roles in ER protein import and ERAD

**DOI:** 10.1101/260174

**Authors:** Francesco Elia, Thomas Tretter, Karin Römisch

**Affiliations:** Department of Microbiology, Faculty of Natural Sciences and Technology VIII, Saarland University, 66123 Saarbruecken, Germany

## Abstract

Sec61p is the channel-forming subunit of the heterotrimeric Sec61 complex that mediates co-translational protein import into the endoplasmic reticulum (ER). In yeast, proteins can also be post-translationally translocated by the hetero-heptameric Sec complex, composed of the Sec61 and the Sec63 complexes. The Sec61 channel is also a candidate for the dislocation channel for misfolded proteins from the ER to the cytosol during ER-associated degradation (ERAD). The structure of the Sec61 complex is highly conserved, but the roles of its N-terminal acetylation and its amphipathic N-terminal helix are unknown so far. To gain insight into the function of the Sec61p N-terminus, we mutated its N-acetylation site, deleted its amphipathic helix, or both the helix and the N-acetylation site. Mutation of the N-acetylation site on its own had no effect on protein import into the ER in intact cells, but resulted in an ERAD defect. Yeast expressing *sec61* without the N-terminal amphipathic helix displayed severe growth defects and had profound defects in post-translational protein import into the ER. Nevertheless the formation of the hetero-heptameric Sec complex was not affected. Instead, the lack of the N-terminal amphipathic helix compromised the integrity of the heterotrimeric Sec61 complex. We conclude that the N-terminal helix of Sec61p is required for post-translational protein import into the ER and Sec61 complex stability, whereas N-terminal acetylation of Sec61p plays a role in ERAD.

## Introduction

Secretory proteins and organelle proteins of the secretory pathway are translocated into the endoplasmic reticulum (ER) during biogenesis [1]. In the ER lumen, imported proteins have to acquire a functional conformation before their delivery to specific cellular destinations via the secretory pathway [2]. Proteins that fail to fold in the ER are retrotranslocated to the cytosol in order to be degraded by proteasomes, a process known as ER-associated degradation (ERAD) [2, 3]. Transport of newly synthesized proteins across the ER membrane can occur either co- or post-translationally [4]. Both modes of translocation require the heterotrimeric Sec61 channel, which consists of three proteins, Sec61p, Sbh1p, and Sss1p in yeast (Sec61α, β, γ in mammals) [5]. The Sec61 complex is sufficient to mediate co-translational import on its own, while it associates with the heterotetrameric Sec63 complex (Sec62p, Sec63p, Sec71p, Sec72p) for post-translational protein import into the yeast ER [5]. Post-translational import generally occurs for soluble proteins that carry only mildly hydrophobic signal sequences, whereas membrane proteins use the signal recognition particle (SRP)-mediated cotranslational pathway [6]. The Sec61 complex is also a candidate channel for the dislocation of ERAD substrates to the cytosol [2, 3, 7, 8].

Sec61p is the channel-forming subunit of the Sec61 complex [9, 10]. The protein is characterized by a compact bundle of 10 transmembrane helices spanning the ER membrane with both termini in the cytoplasm [9, 10]. The two symmetrical halves of Sec61p form an aqueous pore in the ER membrane and a lateral gate facing the lipid bilayer [11]. Sbh1p and Sss1p are tail-anchored membrane proteins with single transmembrane spans [9]. Two evolutionarily conserved large loops of Sec61p, L6 and L8, protruding from the cytoplasmic side of the ER membrane are involved in ribosome binding during co-translational import into the ER [12]. The cytosolic C-terminus of Sec61p has also been shown to contact the ribosome and is functionally important [13, 14]. The cytosolic face of the Sec61 channel also interacts with proteasomes in an ATP-dependent manner [15]. Proteasomes bind the Sec61 channel via the AAA-ATPases of the 19S regulatory particle and *in vitro* compete with ribosomes for ER membrane binding [16]. The AAA-ATPase Cdc48p, involved in the delivery of both misfolded ERAD substrates and partially translocated proteins to the proteasome, can also bind to the Sec61 channel [2, 17]. The specific cytosolic domains of the Sec61 channel responsible for the interaction with AAA-ATPases, however, still remain to be determined [16, 18]. The Sec61 complex also interacts with other transmembrane protein complexes via its small subunits: the mammalian orthologue of Sbh1p, Sec61β, mediates interaction with the signal peptidase complex, and its yeast homologue, Sbh2p, binds to the SRP receptor [19, 20]. Sbh1p and Sss1p also make contact with the oligosaccharyl transferase complex [21, 22].

Despite the fact that Sec61p structure and function have been extensively characterized, the role of its N-terminus is still unknown. The N-terminal region of Sec61p is likely to be functionally important, given that a 6-histidine tag at the Sec61p N-terminus in combination with point mutations elsewhere in the protein interferes with import, but the phenotype is less severe in the absence of the tag [23]. The N-terminus of Sec61p is oriented towards the cytosol and residues 3-21 have the potential to form an amphipathic α-helix [24]. Together with the Sbh1p N-terminus, the Sec61p N-terminus is exposed at one side of the transmembrane helix bundle forming the transmembrane channel and thus poised to make contact with other proteins [14]. In addition, the starting methionine of Sec61p is cleaved and the serine at position 2 acetylated by the NatA complex [25]. N-terminal acetylation occurs co-translationally, and in yeast about 50% of proteins are N-terminally acetylated, but the significance of this modification is only beginning to be understood [26]. Roles for N-terminal acetylation include protein stability, interaction ability, and subcellular targeting [27].

To gain insight into the function of the Sec61p N-terminus, we investigated the effects of altering the sequence context or position of the N-acetylation site by changing the serine at position 2 to tyrosine (*sec61S2Y*) or deleting N-terminal residues 4-22 forming the amphipathic helix (*sec61ΔH1*). In addition, we generated a mutant lacking both the N-terminal acetylation site and the N-terminal residues 4-22 (*sec61ΔN21*). We investigated the effects of these mutations on co- and post-translational import into the ER and ERAD. Furthermore, we asked whether the deletion of the N-terminal helix affected heptameric Sec complex formation and integrity of Sec61 complexes.

## Methods

### Yeast strains

The *sec61-32* point mutant [28] had been previously cloned into the yeast plasmid pRS315 [29]. The *sec61S2Y* mutant was obtained by site-directed mutagenesis of a pRS315 plasmid carrying the *SEC61* gene using the QuikChange kit (Agilent); the second codon of the *SEC61* gene was mutated from TCC to TAC, resulting in a serine to tyrosine amino acid substitution. *sec61ΔH1* and *sec61ΔN21* were obtained by PCR-mediated DNA deletion of a pRS315 plasmid carrying the *SEC61* gene [30]; deletions of the N-terminal residues 4-22 and 2-22 of Sec61p, respectively, were confirmed by DNA sequencing. Plasmids were individually transformed into the KRY461 strain (*sec61::HIS3 ade2-1 leu2-3,112 trp1-1 prc1-1 his3-11, 15 ura3-1 [pGALSEC61-URA3])*, selected on minimal medium without leucine at 30°C, and subsequently selected on minimal medium without leucine at 30°C in the presence of 1 g/l 5- fluoroorotic acid (Sigma).

### Growth conditions

*S. cerevisiae* cells were grown at 30°C in YPD with continuous shaking at 200 rpm or on YPD plates at 30°C. To test temperature sensitivity, 10-fold serial dilutions were prepared and 5 μl of each dilution containing 10^4^-10 cells were dropped onto YPD plates and incubated for 5 (20°C), 3.5 (*sec61ΔH1, sec61ΔN21* and corresponding wildtype; 24°C, 30°C, 37°C), or 3 days (*sec61S2Y, sec61-32* and corresponding wildtype; 30°C, 37°C). To test tunicamycin (Tm) sensitivity, serial dilutions were prepared and 5 of each dilution containing 10^4^-10 cells were dropped onto YPD (±0.25 μg/ml Tm, ±0.5 μg/ml Tm) plates. Plates were incubated at the indicated temperatures for 3.5 (*sec61ΔH1, sec61ΔN21* and corresponding wildtype) or 3 days (*sec61S2Y, sec61-32* and corresponding wildtype).

### β-galactosidase assay

KRY461, *sec61ΔH1, sec61ΔN21* and *sec61S2Y* were transformed with the plasmids pJC31, a pRS314 plasmid carrying the UPRE-LacZ reporter construct, and the control plasmid pJC30, a pRS314 plasmid carrying the LacZ gene [31]. Cells were grown in synthetic complete medium without tryptophan to an OD_600_ of approximately 0.5 and aliquots of 1 ml were harvested by centrifugation and resuspended in 1 ml of Z buffer (60 mM Na_2_HPO_4_, 40 mM NaH_2_PO_4_, 10 mM KCl, 1 mM MgSO_4_, 0.27% β-mercaptoethanol). Subsequently, 100 μl of chloroform and 50 μl of 0.1% SDS were added to each sample; after 10 s of vortexing, samples were pre-incubated for 5 min at 28°C in a water bath, and reactions were induced with 200 of 4 mg/ml 2-nitrophenyl-β-D-galactopyranoside Z buffer. After 20 min of incubation at 28°C, reactions were stopped by adding 500 μl of 1 M Na_2_CO_3_, samples centrifuged, and supernatants analyzed photometrically at 420 nm to calculate β-galactosidase units.

### Western blot analysis

Proteins were resolved by gel electrophoresis using NuPAGE Novex 4-12% Bis-Tris Protein Gels (Thermo Fisher Scientific). Proteins were transferred to nitrocellulose and bands detected by immunoblotting with specific rabbit polyclonal antisera. Antibodies against Sec61p, Sbh1p, CPY [29], Rpn12 [16], and preproalpha factor (ppαF) [29] had been previously raised in our laboratory and were used at a dilution of 1:2000; antibodies against Sss1p and Sec63p were kindly provided by Randy Schekman and used at a dilution of 1:2500. Signals were developed by enhanced chemiluminescence using the SuperSignal West Dura Extended Duration Substrate (Thermo Fisher Scientific) according to the manufacturer’s instructions.

### Cycloheximide chase

Cells expressing chromosomal CPY* [32] were grown overnight in YPD to an OD_600_ of 1 and treated with 200 μg/ml of cycloheximide (t = 0). An equal amount of cells (1 OD_600_) was taken at the indicated time points, washed with sterile water, resuspended in 50 μl of 2x SDS-buffer and lysed with glass beads in a Mini-Beadbeater-16 (BioSpec Products, Bartlesville, OK, US; two 1 min disruption cycles at 4°C with 1 min of incubation at 4°C between cycles). Lysates were heated for 10 min at 65°C and analyzed by Western blotting as described above.

### Pulse-labeling and immunoprécipitation

Cells were grown overnight in YPD to an OD_600_ of 0.5-1 and washed 2 times with labeling medium (5% glucose or galactose, 0.67% yeast nitrogen base without amino acids and ammonium sulfate) supplemented with auxotrophy-complementing amino acids and concentrated to 6 OD/ml. Each 250 cell suspension was preincubated at 30°C under shaking at 600 rpm for 15 min, and subsequently labeled for 5 min adding 55 μCi of EXPRE^35^S^35^S Protein Labeling Mix (PerkinElmer). Labeling was stopped by adding 750 μl of ice-cold Tris-azide (20 mM Tris, pH 7.5, 20 mM sodium azide); cells were subsequently washed with 1 ml of Tris-azide and incubated for 10 min at room temperature in resuspension buffer (100 mM Tris, pH 9.4, 10 mM DTT, 20 mM sodium azide). After centrifugation, cells were lysed with glassbeads in 150 μl of lysis buffer (20 mM Tris, pH 7,5, 2% SDS, 1 mM DTT, 1 mM PMSF) and the lysate denatured for 10 min at 65°C. Glassbeads were washed and the collected supernatant was used for immunoprecipitation with 60 μl of a 20% suspension of Protein A Sepharose beads CL-4B (GE Healthcare) and 10 μl of polyclonal rabbit antisera against CPY or DPAPB in IP buffer (15 mM Tris, pH 7.5, 150 mM NaCl, 1% Triton X-100, 0.1% SDS, 2 mM sodium azide) for 2 h at 4°C. DPAPB antibodies were kindly provided by Tom Stevens. Precipitates were washed in IP buffer, urea wash (100 mM Tris, pH 7.5, 2 M urea, 200 mM NaCl, 1% Triton X-100, 2 mM sodium azide), Con A wash (20 mM Tris, pH 7.5, 500 mM NaCl, 1% Triton X-100, 2 mM sodium azide), Tris-NaCl wash (10 mM Tris, pH 7.5, 50 mM NaCl, 2mM sodium azide), heated for 10 min in 2x SDS-buffer and resolved by gel electrophoresis. Signals were detected by autoradiography using a Typhoon Trio imager (GE Healthcare).

### In vitro post-translational translocation assay

Microsomes were prepared as described in Pilon et al. (1997) [28]. ppαF was translated and ^35^S-methionine-labeled *in vitro* using the Rabbit Reticulocyte Lysate System (Promega) according to the manufacturer’s instructions (2 μg RNA per 50 μl reaction) and translocated into wildtype and mutant microsomes at 20°C for the indicated times. Individual reactions were set up as follows: 3 μl of B88 (20 mM HEPES-KOH, pH 6.8, 250 mM sorbitol, 150 mM KOAc, 5 mM Mg(OAc)_2_), 1 μl of 10x ATP mix (10 mM ATP, 400 mM creatine phosphate, 2 mg/ml creatine kinase in B88), 5 of translation reaction product, 0.6 eq of microsomes (1 μl). To investigate translocation in the presence of limiting amounts of membranes adjusting for equal amounts of Sec61p, 0.3 eq of wildtype and 0.85 eq of *sec61S2Y* microsomes were used for the experiment. After incubation, samples were resolved by gel electrophoresis and signals acquired by autoradiography. Quantitation was performed using the ImageQuant TL software (GE Healthcare).

### Fractionation of Sec complex

Fractionation of Sec complex was performed as described in Pilon et al. (1998) [23]. Briefly, microsomes (50 eq) were resuspended on ice in 100 μl of solubilization buffer (50 mM HEPES-KOH, pH 7.4, 400 mM KOAc, 5 mM MgAc, 10% glycerol, 0.05% β-mercaptoethanol) containing the cOmplete Mini EDTA-free Protease Inhibitor (PI) Cocktail (Roche) and 0.1 mM PMSF; subsequently, 400 μl of solubilization buffer containing 3.75% digitonin were added, resulting in a final 3% digitonin concentration, and samples were incubated on ice for 30 min and subsequently centrifuged at 110,000 g in a TLA 100.3 (Beckman Instruments, Palo Alto, CA, US) rotor for 30 min at 4°C. The Sec63 complex was precipitated from the resulting supernatant at 4°C for 1 h with 100 μl of Concanavalin A (Con A) Sepharose 4B (GE Healthcare). Remaining beads were cleared from the supernatant by centrifugation to obtain the free fraction. Con A beads were washed twice with equilibration buffer (1% digitonin, 50 mM HEPES-KOH pH 7.4, 10% glycerol, 0.05% β-mercaptoethanol, 0.1 mM PMSF and PIs) and glycoproteins were subsequently released in 2x SDS-buffer for 10 min at 65°C. To obtain the ribosome-associated membrane protein (RAMP) fraction, the pellet from the 110,000 g spin was dissolved in 100 of 50 mM HEPES-KOH (pH 7.8), 1 M KAc, 17.5 mM MgAc, 2.5% digitonin, 1 mM puromycin, 0.2 mM GTP, 5 mM dithiothreitol, 0.1 mM PMSF containing PIs. After 30 min on ice and 30 min at 30°C, RAMPs including the Sec61 complex were recovered from the supernatant after centrifugation at 100,000 g for 30 min at 4°C. Equal amounts of each fraction were analyzed by gel electrophoresis on NuPAGE Novex 4-12% Bis-Tris Protein Gels (Thermo Fisher Scientific) and immunoblotting with the indicated antibodies.

### Sucrose gradient centrifugation

Sucrose gradients were prepared in 13 x 51 mm polycarbonate centrifuge tubes (Beckman) using 1 ml of 15%, 10%, 5% and 0% sucrose in 50 mM HEPES-KOH, pH 7.5, 500 mM potassium acetate, 1 mM EDTA, 0.1% Triton X-100, 0.05% β-mercaptoethanol, 1 mM PMSF containing PIs. Microsomes (50 eq) were centrifuged and the pellet was resuspended in 100 μl of solubilization buffer (50 mM HEPES-KOH, pH 7.5, 500 mM KOAc, 1% Triton X-100, 10 mM EDTA, 0.05% β-mercaptoethanol, 1 mM PMSF and 5x PIs) and incubated for 15 min on ice. Solubilized microsomes were loaded onto 0-15% sucrose gradients and ultracentrifuged (200,000 g, 4°C, 16 h) in a TLA 100.3 rotor. After centrifugation, 13 fractions were collected from top to bottom of the gradients, precipitated with TCA, and analyzed by gel electrophoresis and immunoblotting with the indicated antibodies.

## Results

Viability and ER stress induction are affected in *sec61* N-terminal mutants

To investigate the function of the Sec61p N-terminus, we used *sec61* mutants carrying a deletion of the N-terminal helix but preserving the N-terminal acetylation site in its original sequence context, *sec61ΔH1*, lacking both the N-terminal acetylation site and the N-terminal helix, *sec61ΔN21*, or carrying a mutation in the N-acetylation acceptor site, *sec61S2Y* (Fig. 1A). The serine to tyrosine substitution in position 2 results in a non-cleavable and acetylated initiator methionine, and thus in a substantially more bulky N-terminus than in the wildtype protein [33, 34]. Mutants were introduced into a strain bearing a chromosomal deletion of *SEC61* by plasmid shuffling [29]. All *sec61* mutant proteins were stable, and expressed at wildtype level (*sec61ΔH1)* or approximately 40% of wildtype (*sec61S2Y* and *sec61ΔN21*, data not shown). We first tested the effects of these mutations on cell growth at various temperatures and in the presence or absence of tunicamycin, which interferes with N-linked glycosylation in the ER and hence protein folding [35]. Translocation-defective *sec61* mutants frequently accumulate misfolded proteins in the ER and are therefore tunicamycin-sensitive [29]. The *sec61S2Y* mutant, together with the ERAD-defective cold-sensitive *sec61- 32* strain as a control, was grown at 20°C, 30°C, and 37°C without tunicamycin, and at 30°C in the presence of 0.25 or 0.5 μg/ml tunicamycin (Fig. 1B). The *sec61S2Y* mutant did not grow at 37°C (Fig. 1B, top panel), suggesting a possible role for the acetylation of S2 in Sec61p function. In addition, growth of the strain was affected in the presence of the higher concentration of tunicamycin, indicating perturbations in ER homeostasis. Growth of the *sec61ΔH1* and *sec61ΔN21* mutants was analyzed at 20°C, 30°C, and 37°C, both in the presence and absence of 0.25 μg/ml tunicamycin (Fig. 1C). The strains exhibited severe growth defects under all conditions tested and no growth at all at 24°C in the presence of tunicamycin (Fig. 1C). These results indicate that the N-terminal domain of Sec61p is highly important for function.

**Fig 1.**
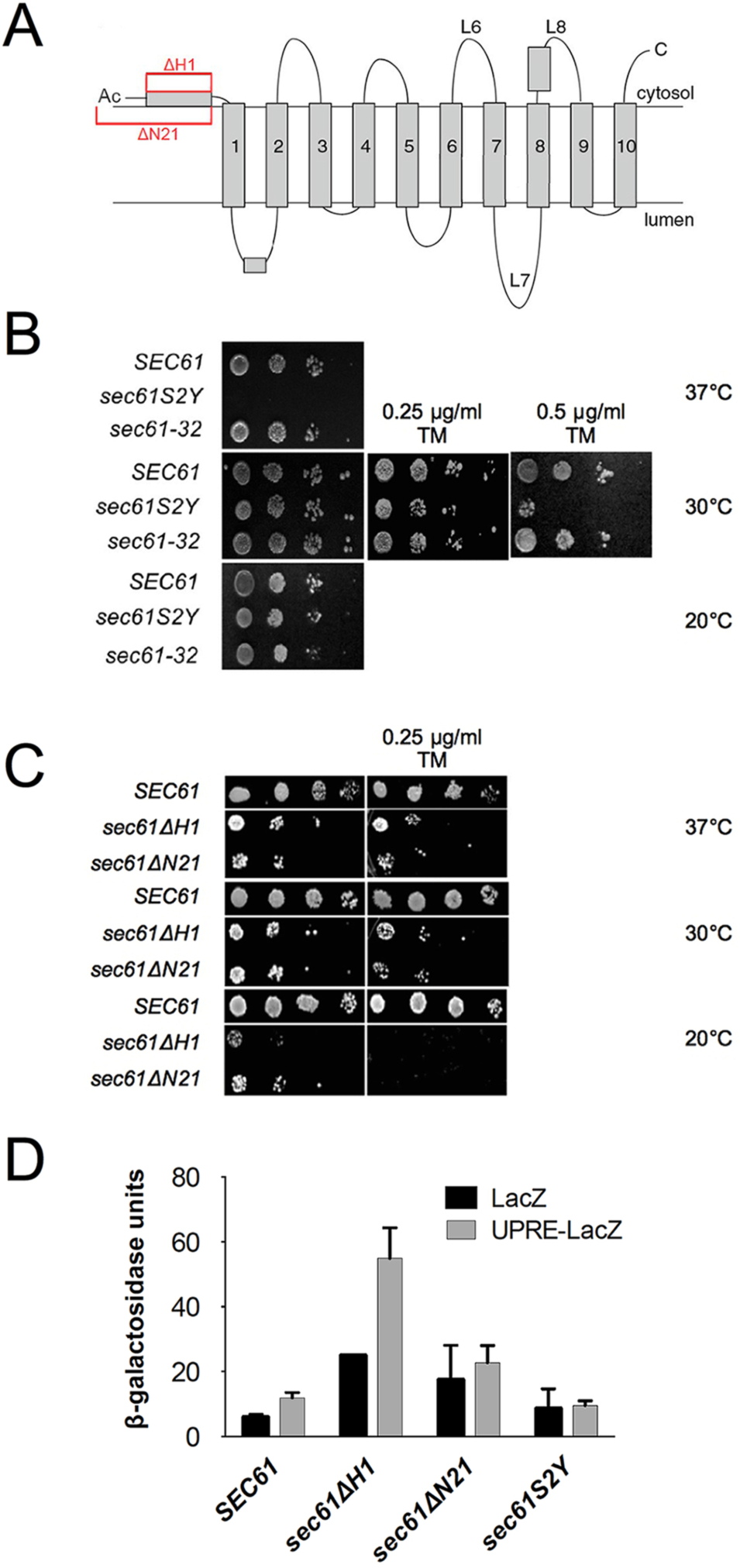
Viability and growth of *sec61* N-terminal mutants under stress conditions. A: Topology of Sec61p. N-terminal mutations characterized in this paper are highlighted in red. B: Temperature- and tunicamycin-sensitivity of *sec61S2Y.* Cells (10^4^-10) expressing *sec61S2Y, sec61-32*, or wildtype *SEC61* were grown at 20°C (6 d), 30°C (without or with 0.25 μg/ml and 0.5 μg/ml tunicamycin) and 37°C (3 d) on YPD plates. C: Temperature- and tunicamycin-sensitivity of *sec61ΔH1* and *sec61 AN21*. Cells (10^4^-10) expressing *sec61ΔH1, sec61 AN21*, or wildtype *SEC61* were grown for 3.5 days on YPD plates without or with 0.25 μg/ml tunicamycin at the indicated temperatures. D: Liquid β-galactosidase assay with *sec61* N-terminal mutants. The *sec61ΔH1, sec61 AN21*, and *sec61S2Y* strains were transformed with the plasmids pJC31 (UPRE-LacZ) and the control plasmid pJC30 (LacZ). Cells were harvested in early exponential phase, lysed, and β-galactosidase activity was analyzed photometrically in duplicate samples. The experiment was performed 2 times with duplicate samples. Error bars represent standard deviation.

To directly investigate UPR induction of the *sec61ΔH1, sec61ΔN21* and *sec61S2Y* strains, we transformed each of the mutants and the corresponding wildtype with plasmids carrying the β-galactosidase gene under the control of a promoter without or with a UPR element [31]. Transformants were grown at 30°C, lysed, and β-galactosidase activity monitored using a chromogenic substrate [31]. The UPR was strongly induced in the *sec61ΔH1* strain, but although the β-galactosidase activity was higher in *sec61ΔN21* than in wildtype cells, this was independent of the UPRE (Fig. 1D). Despite its temperature- and tunicamycin-sensitivity, in the *sec61S2Y* mutant the UPR was not induced (Fig. 1D). Collectively, our data suggest that repositioning of the N-terminal acetylation site in *sec61ΔH1* is more detrimental to ER protein homeostasis than the absence of N-acetylation in the *sec61ΔN21* mutant.

### N-terminal acetylation at S2 of Sec61p plays a role in ERAD

Tunicamycin-sensitivity is often associated with defects in export of misfolded proteins from the ER to the cytosol [36]. We therefore conducted cycloheximide chase experiments to monitor the decline of the steady state levels of the commonly used soluble ERAD substrate CPY* in the *sec61S2Y, sec61ΔH1*, and *sec61ΔN21* strains [32]. We found that the half-life of CPY* increased approximately 2-fold in *sec61S2Y* compared to the wildtype strain (Fig. 2A). In the helix deletion mutants *sec61ΔN21* and especially *sec61ΔH1* CPY* accumulated in the ER during the chase, suggesting that post-translational import of CPY* into the ER was still taking place after protein biosynthesis had been inhibited with cycloheximide (Fig. 2B).

**Fig 2.**
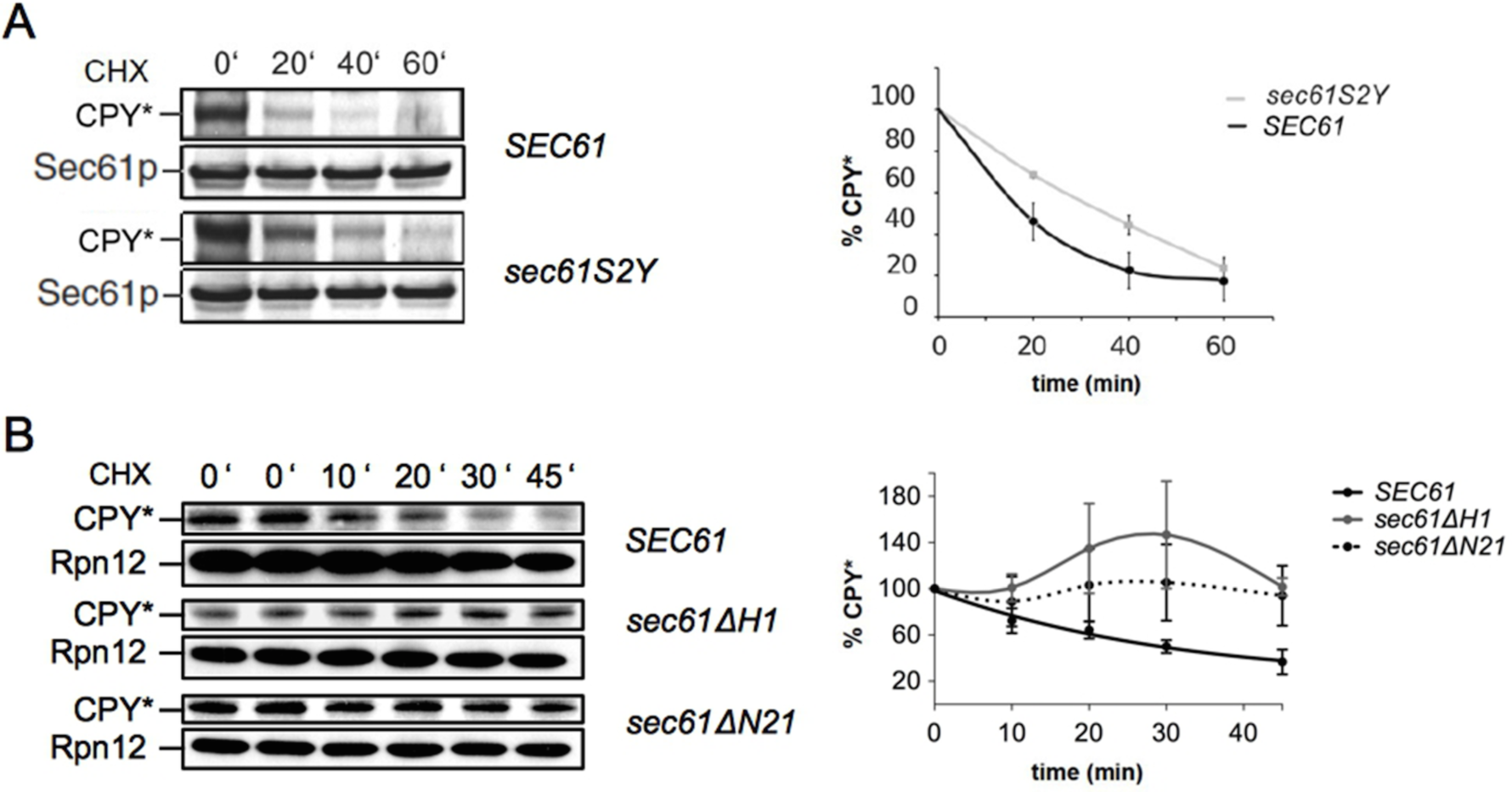
Sec61p N-acetylation at S2 is important for ERAD. ERAD of CPY* was investigated in *sec61S2Y*(A), *sec61ΔH1*, and *sec61ΔN21* (B). Cells were grown at 30°C to an OD_600_ = 1, translation was inhibited by adding 200 μg/ml cycloheximide, extracts were prepared by bead-beating, and samples resolved by SDS-PAGE. The 0' samples were taken in duplicate in (B). Proteins were transferred to nitrocellulose and CPY* was detected with polyclonal rabbit antiserum and enhanced chemiluminescence. Bands were quantified using the ImageQuant TL software. Sec61 (A) and Rpn12 (B) were used as loading controls. Band intensities were quantified relative to loading controls. Curves represent the averages of three independent experiments. Error bars represent standard deviations.

Whether or not ERAD was also affected in these mutants remained unclear, even when we extended the chase to 90 min (data not shown). We conclude that N-acetylation of Sec61p at S2 is required for ERAD of misfolded soluble proteins.

### Deletion of the N-terminal amphipathic helix of Sec61p severely impairs post-translational protein import into the ER

As the results shown in Fig. 2B suggest an ER import defect or an ERAD defect for *sec61ΔH1* and *sec61ΔN21*, we decided to investigate import directly. We first monitored co-translational membrane integration of diaminopeptidase B (DPAPB) in the *sec61S2Y, sec61ΔH1*, and *sec61ΔN21* strains by pulse-labeling with ^35^S-Met/Cys for 5 min, lysing the cells, and immunoprecipitation with specific polyclonal antibodies. We barely detected the cytosolic precursor form of DPAPB (pDPAPB) in *sec61ΔH1* and *sec61ΔN21* (Fig. 3A, right panel), and mutation of the N-terminal acetylation site had no effect at all on pDPAPB import (Fig. 3A, left panel). These data show that none of the investigated N-terminal *sec61* mutations affects co-translational translocation into the ER.

**Fig 3.**
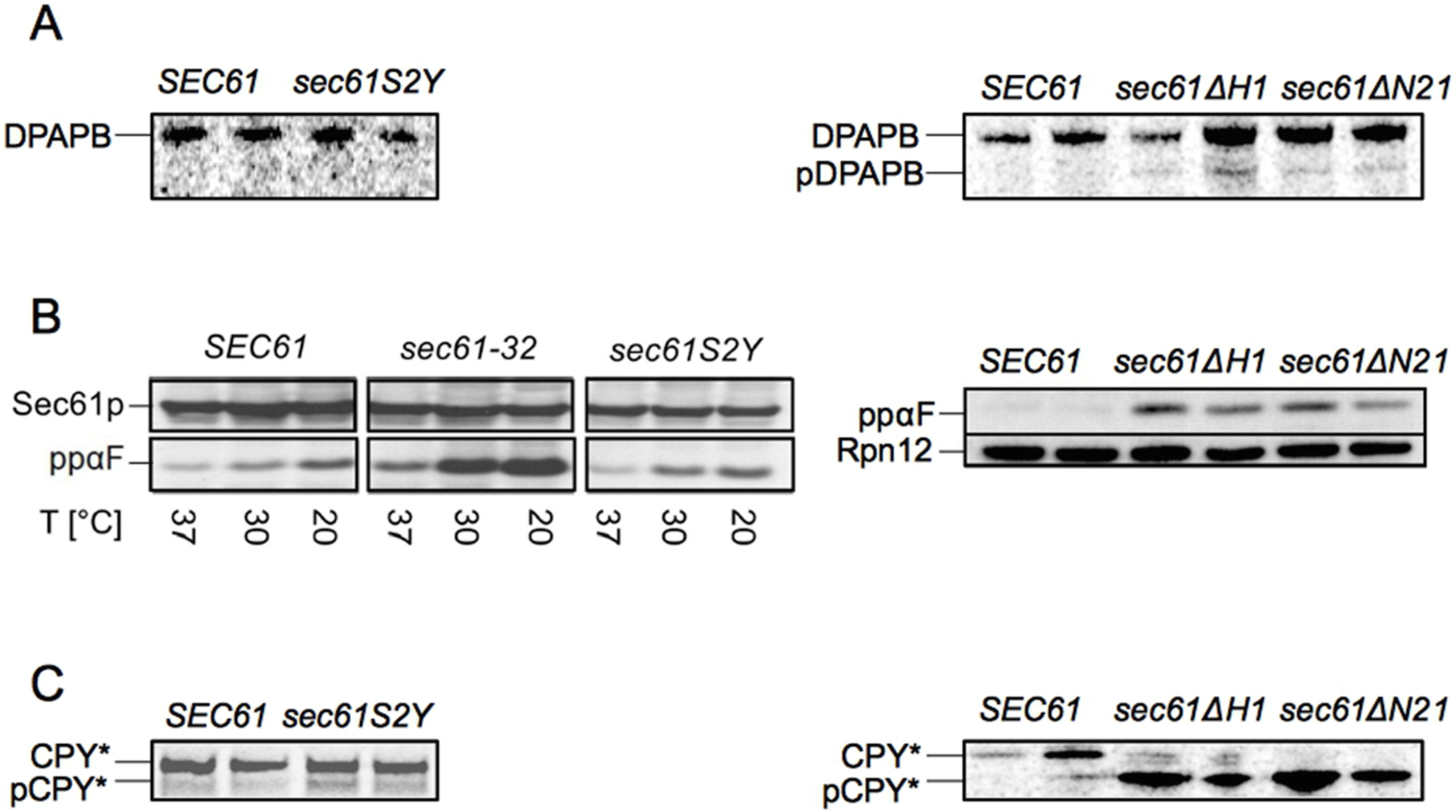
The Sec61p N-terminal helix is important for post-translational protein import into the ER *in vivo.* A: Co-translational ER import of newly synthesized DPAPB in *sec61S2Y*(left panel), *secóJAHJ*, and *sec61ΔN21* (right panel). Cells were grown at 30°C to early exponential phase, labeled with ^35^S-Met/Cys for 5 min and lysed. Cytosolic precursor (pDPAPB) and ER-membrane integrated DPAPB were immunoprecipitated, resolved by SDS-PAGE, and detected by autoradiography. B: Post-translational ER import of ppαF in *sec61S2Y* (left panel), *sec61ΔH1*, and *sec61ΔN21* (right panel). Cells were grown at 30°C to an OD600 = 1 and shifted to the indicated temperatures for 3h, extracts were prepared by bead-beating, and samples resolved by SDS-PAGE. Proteins were transferred to nitrocellulose and the accumulation of cytosolic ppαF was analyzed by immunoblotting. C: Post-translational ER import of newly synthesized pCPY* in *sec61S2Y* (left panel), *sec61ΔH1*, and *sec61ΔN21* strains (right panel). Cells were grown at 30°C to early log phase, labeled with ^35^S-Met/Cys for 5 min and lysed; cytosolic pCPY* and ER-lumenal CPY* were immunoprecipitated, resolved by SDS-PAGE and detected by autoradiography. Duplicate samples were taken for experiments shown in panels A, B (right), and C. Experiments were performed three times.

We next investigated the *sec61S2Y, sec61ΔH1*, and *sec61ΔN21* mutants for post-translational import defects, monitoring the cytoplasmic accumulation of the post-translational import substrate preproalpha factor (ppαF) in cell lysates by Western Blotting with specific polyclonal antisera. Substitution of the N-acetyl acceptor serine in position 2 of Sec61p had no effect on post-translational import of ppαF *in vivo* (Fig. 3B, left panel). In addition, cytosolic accumulation of ppαF in *sec61S2Y* was not increased at 37°C, despite the temperature sensitivity of the *sec61S2Y* mutant (see Fig. 1B). In the cold-sensitive *sec61-32* control strain, however, ppαF levels were increased at 20°C (Fig. 3B, left panel). As shown in Fig. 3B (right panel), post-translational import of ppαF was profoundly affected in both *sec61ΔH1* and *sec61ΔN21.* Strong post-translational import defects were also displayed by the *sec61ΔH1* and *sec61ΔN21* mutants in a pulse-labeling experiment with ^35^S-Met/Cys for 5 min followed by immunoprecipitation with antibodies against carboxypeptidase Y (CPY) to monitor the efficiency of post-translational ER import of newly synthesized proteins (Fig. 3C, right panel). No post-translational import of cytosolic precursor pCPY* could be detected in *sec61ΔN21* cells during the 5 min pulse, and the *sec61ΔH1* mutant only allowed minimal pCPY* import into the ER (Fig. 3C, right panel). In contrast, the *sec61S2Y* substitution led only to a marginal accumulation of pCPY* in the cytosol (Fig. 3C, left panel). Our data suggest that the N-terminal helix of Sec61p, but not its N-terminal acetylation site, is essential for post-translational import into the ER.

### Post-translational protein import into *sec61S2Y, sec61ΔH1* and *sec61ΔN21* microsomes is impaired *in vitro*

As subtle translocation defects can be masked by the abundance of Sec61 channels in intact cells, to further explore a potential impact of the *sec61S2Y* mutation on protein import into the ER, we investigated the ability of *sec61S2Y* microsomes to import ppαF *in vitro* [18, 37]. The ppαF mRNA was translated *in vitro* in the presence of ^35^S-labeled methionine, and the resulting radiolabeled ppαF subsequently incubated at 20°C in the presence of ATP and limiting amounts of wildtype or mutant microsomes for the indicated periods of time (Fig. 4A). *In vitro* the *sec61S2Y* mutation led to a reduction in the import of ppαF into yeast microsomes (Fig. 4A), and this post-translational import defect was more substantial compared to the one found in intact cells (compare Fig. 4A vs. Fig. 3C, left panel). We also attempted to investigate ppαF import into *sec61ΔH1* and *sec61ΔN21* microsomes (Fig. 4B). Even after 2 hours of incubation, however, we detected no glycosylated 3gpαF in the *sec61ΔN21* samples, and only a very limited amount of 3gpαF in the presence of *sec61ΔH1* membranes (Fig. 4B). These results confirm the critical role of the Sec61p N-terminus in post-translational soluble protein import into the ER. Our data suggest a requirement for the Sec61p N-terminal amphipathic helix for post-translational protein import into the ER with a possible contribution by N-acetylation at the N-terminus of the helix.

**Fig 4.**
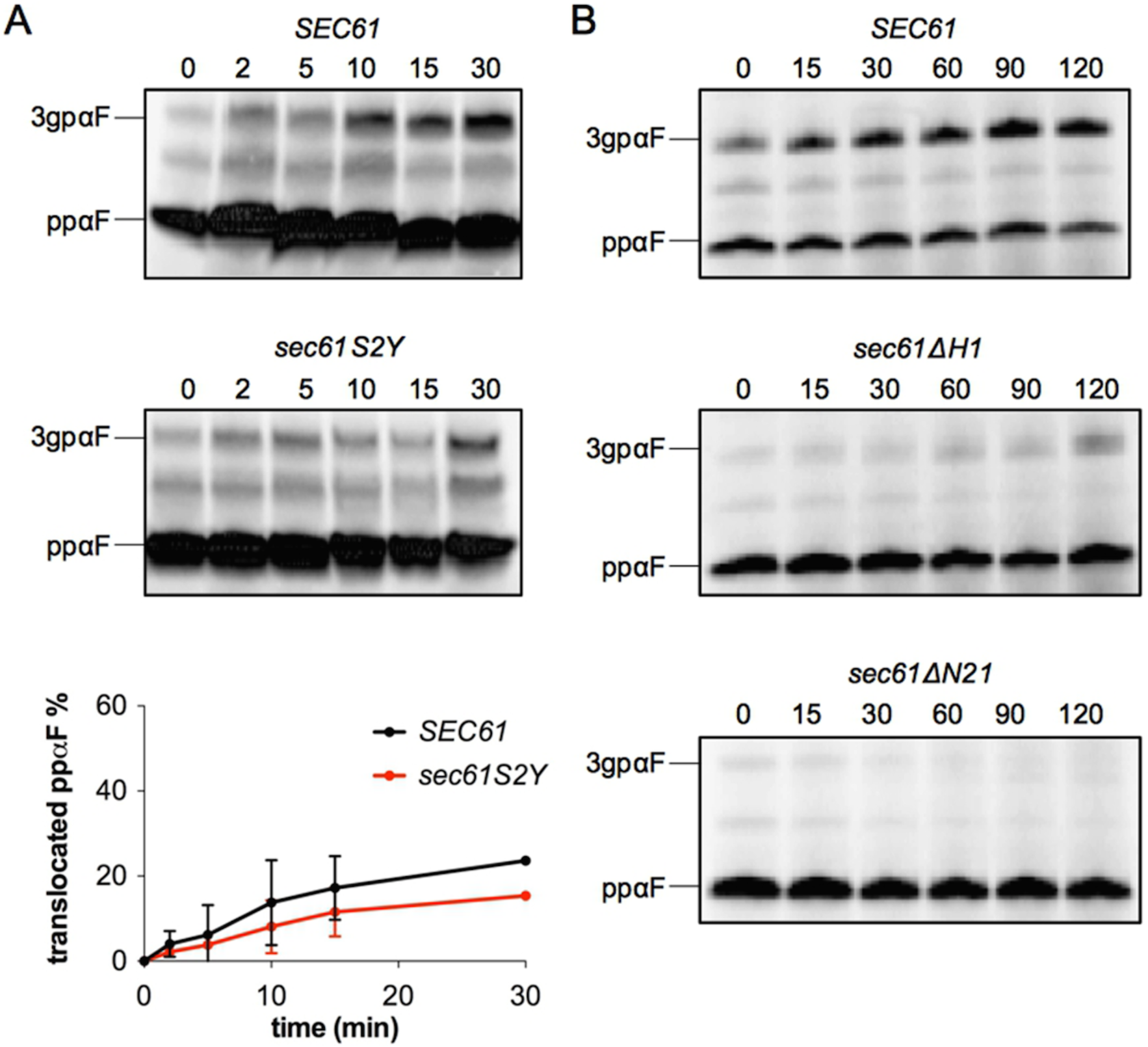
N-acetylation of Sec61p at S2 and its N-terminal helix are required for ER import *in vitro.* *In vitro* post-translational import of ppαF into limiting amounts of *sec61S2Y* (A), *sec61ΔH1*, and *sec61ΔN21* microsomes (B). The mRNA for ppαF was translated in the presence of ^35^S-methionine *in vitro* and radiolabeled ppαF was translocated into wildtype and mutant microsomes at 20°C in the presence of ATP and a regenerating system. At the indicated time points (min), 2x SDS-buffer was added and samples were resolved by SDS- PAGE and cytosolic ppαF and translocated, signal-cleaved glycosylated 3gpαF detected by autoradiography. Bands were quantified using the ImageQuant TL software. Experiments were performed four times. The graph in A represents the average of two independent experiments. Error bars represent standard deviation.

### The Sec61 complex is unstable in the absence of the N-terminal helix

To test whether the severe defects in post-translational import found for *sec61ΔH1* and *sec61ΔN21* both *in vivo* and *in vitro* (Fig. 3 & Fig. 4) were due to an impaired formation of heptameric Sec complexes, we solubilized wildtype, *sec61ΔH1*, and *sec61ΔN21* microsomes in digitonin, ultracentrifuged lysates, and precipitated Sec complexes from the supernatant using Concanavalin A beads, which bind to the N-glycans of Sec71p [23]. The solubility of Sec61ΔH1p and Sec61ΔN21p was dramatically reduced compared to wildtype, suggesting that Sec61p without its N-terminal helix is prone to aggregation (Fig. 5A, Sol. fractions). Although we could only solubilize small amounts of Sec61ΔH1p and Sec61ΔN21p, we were able to detect amounts of both variants in the Con A fractions although the ratios of Sec61ΔH1p and Sec61ΔN21p to Sec63p in the ConA fractions were lower compared to wildtype (Fig. 5A, ConA fractions). In contrast, we observed a dramatic loss of both mutant *sec61* protein variants from the ribosome-associated membrane protein (RAMP) fractions (Fig. 5A). We found Sec63p in comparable amounts in the Con-A and RAMP fractions of wildtype, *sec61ΔH1*, and *sec61ΔN21* membranes, but there was more Sec63p in the “Free” fraction in the mutants compared to wildtype (Fig. 5A, bottom panels). Our data suggest that despite the strong post-translational import defects shown by both the N-terminal helix deletion mutants, their heptameric Sec complexes are largely intact.

**Fig 5.**
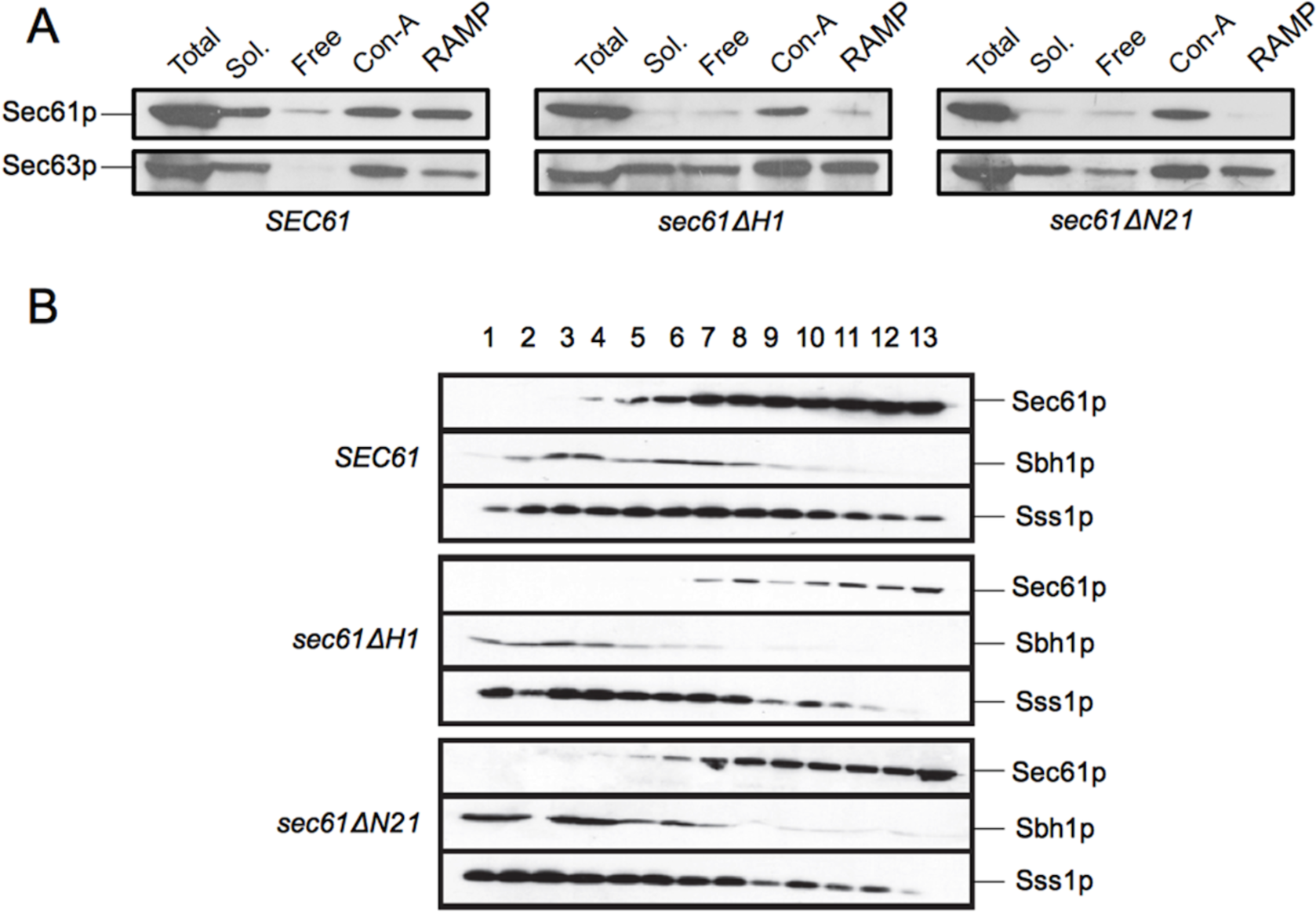
The N-terminal helix of Sec61p is required for Sec61 complex stability. A: Stability of heptameric Sec complexes in *sec61ΔH1* and *sec61ΔN21* membranes. Wildtype, *sec61ΔH1*, and *sec61ΔN21* microsomes were solubilized in solubilization buffer (see Methods) containing 3% digitonin and lysates centrifuged at 110,000 g to sediment ribosome-bound Sec61 complexes (ribosome-associated membrane proteins, RAMP). From the supernatants, heptameric Sec complexes were precipitated with Concanavalin A Sepharose (Con A). Sec61p and Sec63p not bound to either Con A or ribosomes were detected in the “Free” fraction. In all fractions, Sec61p and Sec63p were detected by immunoblotting with polyclonal antisera. Loss of Sec61p from the *sec61ΔH1* and *sec61ΔN21* Soluble fractions is likely due to the formation of SDS-resistant aggregates in digitonin. B: Stability of Sec61 complexes in *sec61ΔH1* and *sec61ΔN21* strains. Microsomes were solubilized in 1% Triton X-100 and layered onto a 0-15% sucrose gradient. After centrifugation at 200,000 g for 16 h, fractions were collected from top to bottom, and proteins resolved by SDS-PAGE. Sec61p, Sbh1p, and Sss1p were detected by immunoblotting. For both A and B the experiment was performed 4 times.

By contrast, either the stability of the Sec61 complex or its interaction with ribosomes seemed to be compromised in the absence of the N-terminal helix, as indicated by the reduced amount of mutant Sec61p in the RAMP fractions (Fig. 5A, top panels). Next, we therefore investigated the stabilities of the *sec61ΔH1* and *sec61ΔN21* trimeric complexes directly. We solubilized wildtype, *sec61ΔH1*, and *sec61ΔN21* microsomes in 1% Triton X-100 and centrifuged solubilized Sec61 complexes through a shallow 0-15% sucrose gradient [38]. Fractions were collected from top to bottom, proteins precipitated with trichloroacetic acid, resolved by SDS-PAGE, and Sec61 complex subunits detected in each fraction by blotting with antibodies against the C-termini of Sec61p, Sbh1p, and Sss1p. We observed a distinct loss of colocalization of Sec61p, Sbh1p, and Sss1p in gradient fractions from both *sec61ΔH1* and *sec61ΔN21* complexes (Fig. 5B). A reduced number of fractions contained all three proteins for the two deletion mutants (wildtype: fractions 4-10; *sec61ΔH1:* fraction 7; *sec61 AN21:* fractions 5-7). The destabilization of trimeric Sec61 complexes was more pronounced in *sec61ΔH1* microsomes compared to *sec61ΔN21* microsomes. In contrast, solubilization and fractionation of microsomes derived from *sec61S2Y* cells showed that the interaction of Sec61S2Yp with Sbh1p and Sss1p was not significantly altered (data not shown). Our data suggest that the N-terminal amphipathic helix of Sec61p is required for stability of the Sec61 complex.

## Discussion

The Sec61 complex mediates protein import into the ER, and is also a candidate channel for the dislocation of ERAD substrates [3, 5]. To investigate the function of the Sec61p N-terminus in these processes in yeast, we characterized a set of *sec61* N-terminal mutants. We have shown here that N-acetylation of Sec61p at S2 is important for ERAD (Fig. 2A), and may contribute to post-translational import into the ER (Fig. 4A), whereas its N-terminal amphipathic helix is essential for post-translational import into the ER and is required for stability of the Sec61 complex (Fig. 3C, Fig. 5B).

### Role of Sec61p N-acetylation

We investigated Sec61p function in the *sec61S2Y* mutant, in which Sec61p is not acetylated at its canonical NatA consensus serine at position 2 after methionine cleavage, but rather at the uncleaved initiator methionine, and in two new *sec61* mutants: one carrying a deletion of the N-terminal helix but preserving the N-terminal acetylation site in its original sequence context, *sec61ΔH1*, and one lacking both the N-terminal acetylation site and the N-terminal helix, *sec61ΔN21* (Fig. 1A). The mutant strains stably expressed Sec61p at wildtype level (*sec61ΔH1)* or approximately 40% of wildtype (*sec61S2Y* and *sec61 ΔN21*, data not shown). In a GAL shut-off experiment ER translocation defects only occurred when Sec61p levels fell below 20% of wildtype, and we observed no effect of *sec61S2Y* on cotranslational and only a marginal effect on posttranslational import *in vivo*, confirming that Sec61p was not limiting in our N-terminal *sec61* mutant cells (KR, unpublished; Fig. 3A, 3B, 3C, left panels). N-acetylation at the initiator methionine can target soluble proteins for degradation by the Ac/N-end rule pathway, but since Sec61S2Yp was stable in a cycloheximide chase over 3 h this does not seem to be the case for our mutant protein (data not shown) [39]. Since mutation of the N-terminal acetylation site did not influence the stability of Sec61p, but resulted in lower expression levels, our data suggest that N-acetylation at S2 may play a role in biosynthesis of Sec61p.

Although *sec61S2Y* cells were temperature-sensitive, incubation at higher temperature did not affect co- or post-translational import (data not shown; Fig. 3B, left panel). In *in vitro* experiments with limiting amounts of microsomes and adjusting for equal amounts of Sec61p and Sec61S2Yp, however, we detected a reduced post-translational import of ppαF into *sec61S2Y* microsomes (compare Fig. 4A vs. Fig. 3C, left panel). This difference between the *sec61S2Y* effect in intact cells and in the cell-free assay might be due to an increase in mutation-associated instability of Sec61p as a consequence of the microsome preparation procedure. N-terminal acetylation has been shown to increase N-terminal helicity and affinity for physiological membranes, and although N-acetylation in the *sec61S2Y* mutant is preserved, the helix stabilizing effect is likely lost due to the increased bulk of the N-terminus generated by the presence of both the initiator methionine and the tyrosine at position 2 [40]. Thus, the S2Y mutation will lead to fraying of the N-terminal helix, which is essential for post-translational import (below).

Fig. 2A shows that CPY* retrotranslocation to the cytosol is considerably delayed in the *sec61S2Y* mutant. Since the mutant has no protein import defect *in vivo* (Fig. 3), this effect is likely direct. The defect in ERAD of CPY* was not exacerbated in *sec61S2Y* cells at higher temperature, thus the temperature-sensitivity of the mutant remains unexplained (data not shown). Since protein-protein interactions are often dependent on N-terminal acetylation in a specific sequence context, our data may indicate that Sec61p N-acetylation at S2 is required for interaction with components of the ERAD machinery like the Hrd1 ligase, or the Cdc48 complex [16, 17, 33, 41].

The *sec61ΔH1* mutant was the only N-terminal *sec61* mutant that displayed a significant UPR induction, although it preserves the N-acetylation site (Fig. 1D). Since the mutant protein is stable, this is unlikely to be a result of Sec61ΔH1p eliciting the UPR itself as a misfolded protein. The N-acetylation of Sec61ΔH1p, however, may recruit an interacting factor to its N-terminus, which in the mutant protein due to the absence of the helix ends up in the wrong position and further interferes with function of Sec61ΔH1p [41].

### Role of the Sec61p N-terminal amphipathic helix

Growth of the *sec61ΔH1* and *sec61ΔN21* mutants was compromised in all conditions tested, suggesting an important role for the N-terminal amphipathic helix in Sec61p function (Fig. 1C). The growth defect was caused by a strong post-translational import defect (Fig. 3B, 3C, right panels), which - due to the overlap between import and export during the chase - made it difficult to evaluate possible ERAD defects in the helix deletion mutants (Fig. 2B). In addition, we found that all *sec61* N-terminal mutants including *sec61S2Y* were synthetically lethal with an inefficiently translocating post-translational ER import substrate that clogs the channel, suggesting that the Sec61p N-terminus enhances efficiency of post-translational protein import into the ER (data not shown) [42].

A possible explanation for the almost exclusive post-translational import defects exhibited by the *sec61ΔH1* and *sec61ΔN21* mutants was that the interaction of Sec61p with the Sec63 complex required for post-translational import is compromised in the absence of the N-terminal helix. We examined this by Con A precipitation of solubilized Sec complexes (Fig. 5A), but to our surprise found no reduction in Sec complex formation in the mutants (Fig. 5A). Instead, we found a dramatic loss of Sec61p from the RAMP fractions of *sec61ΔH1* and *sec61ΔN21* membranes, suggesting reduced affinity of mutant Sec61 complexes for ribosomes or reduced Sec61 complex stability (Fig. 5A).

Gradient fractionation experiments with solubilized Sec61 complexes corroborated the hypothesis that the N-terminal helix of Sec61p is important for Sec61 complex stability (Fig. 5B). Membranes were solubilized with Triton X-100 because Sec61 complexes are only marginally stable in this detergent, thus even minor perturbations result in dissociation of the subunits [38]. The distribution of Sec61 complex subunits in the gradient fractions suggests that the Sec61p N-terminal helix is required for Sec61 complex stability, and that interaction of Sbh1p with Sec61p is more strongly dependent on the Sec61p N-terminal helix than that of Sss1p with Sec61p, as Sbh1p colocalized with Sec61ΔH1p in only 1 and with Sec61ΔN21p in only 3 fractions, respectively (Fig. 5B). Sbh1p consists of a single transmembrane helix that is close to transmembrane domain 4 of Sec61p in the crystal structure, and its cytosolic domain is physically close to the cytosolic N-terminus of Sec61p [10, 43]. Upon solubilization, the interaction between the cytosolic domains of Sec61p and Sbh1p might become more important, and the dissociation of Sbh1p from Sec61ΔH1p and Sec61ΔN21p after Triton X-100 solubilization suggests that the Sec61p N-terminal helix might be the cytosolic Sbh1p interacting partner (Fig. 5B). The growth defects of *sec61ΔH1* and *sec61ΔN21* cells were, however, not rescued upon overexpression of *SBH1*, suggesting that Sbh1p interaction is not limiting for mutant channel stability (data not shown). Sss1p makes contact to Sec61p at multiple sites, including the transmembrane helices H6, H7, and H8 [44]. The reduced interaction of Sss1p with Sec61p lacking its N-terminal helix (see Fig. 5B; wildtype: fractions 4-13; *sec61ΔH1:* fractions 7-12; *sec61ΔN21:* fractions 7-12) might therefore be caused by structural changes in the overall conformation of Sec61p that affect some of its interaction sites with Sss1p. Structure of the N-terminal half of Sec61p might be stabilized by the association of the amphipathic N-terminal helix with the cytosolic ER membrane surface and thus coordinated movement of the N-terminal half of Sec61p during channel opening might be compromised if the helix is missing [10]. Given that we observed no DPAPB import defects in *sec61ΔH1* and *sec61ΔN21* cells (Fig. 3A, right panel), the Sec61 complex lacking the N-terminal helix seems to be sufficiently stabilized by the ribosome to function during co-translational protein import into the ER. Our data suggest that, in the absence of the N-terminal helix of Sec61p, the Sec61 complex is unstable, and that its interactions with the Sec63 complex are insufficient to stabilize the channel for post-translational import.

In summary, we have shown here that the Sec61p N-terminus contains two structural elements that are functionally important: N-acetylation at S2 plays a role in ERAD, likely stabilizes the N-terminal helix, and may play a role in Sec61p biogenesis, whereas the Sec61p N-terminal amphipathic helix is essential for post-translational import into the ER and Sec61 complex stability.

## Acknowledgements

Antibodies against Sss1p and Sec63p were kindly provided by Randy Schekman. DPAPB antibodies were kindly provided by Tom Stevens. A first draft of the manuscript was generated by the M.Sc. Infectious Disease Biology class of 2016.

## Author contributions

Conceived and designed the experiments: FE TT KR. Performed the experiments: FE TT. Analyzed the data: FE KR. Contributed reagents/materials/analysis tools: FE KR. Wrote the paper: FE KR.

